# Spatial targeting of the prostaglandin receptor EP2 to very early endosomes co-ordinates PGE2-mediated cAMP signaling in decidualizing human endometrium

**DOI:** 10.1101/2024.03.07.582712

**Authors:** Paul J. Brighton, Abigail R. Walker, Oliver Mann, Chow-Seng Kong, Emma S. Lucas, Pavle Vrljicak, Jan J. Brosens, Aylin C. Hanyaloglu

## Abstract

Decidualization denotes the differentiation of endometrial stromal cells into specialized decidual cells, essential for embryo implantation and pregnancy. The process requires coordinated activation of both progesterone and cAMP signaling pathways, which converge on downstream transcription factors. PGE2 and relaxin, acting respectively through their Gαs-coupled GPCRs EP2 and RXFP1, are putative candidates responsible for generating cAMP in differentiating stromal cells. Here, we show that PGE2 is less efficacious than relaxin in elevating intracellular cAMP levels in primary stromal cells but more effective at driving the expression of canonical decidual genes. Both PGE2- and relaxin-induced cAMP generation involves receptor internalization, but EP2 is endocytosed into very early endosomes (VEEs). Perturbation of the VEE machinery dysregulates PGE2-dependent cAMP profiles and disrupts key decidual signaling pathways, resulting in a disordered differentiation response. We demonstrate that the spatial location of EP2 is essential for coordinated activation of the downstream signaling cascades that govern decidualization.

## Introduction

The human endometrium undergoes profound and dynamic remodeling across the menstrual cycle, with changes marked by cyclical phases of menstrual repair, estrogen-dependent proliferation, progesterone-dependent differentiation and senescence-like tissue break-down and shedding. ^1,2^ In particular, the process of decidualization, denoting the differentiation of endometrial stromal fibroblasts into specialized secretory decidual cells, occurs during the secretory phase of the menstrual cycle after ovulation, concomitant with the secretory transformation of endometrial glands, vascular remodeling, and immune cell recruitment. The process creates a supportive and nutritive environment that is indispensable for embryo implantation, placentation and pregnancy. ^3^ Decidualization is driven by ovarian progesterone which activates the nuclear progesterone receptor (PGR) in stromal cells. However, the process also requires a sustained intracellular increase in the second messenger molecule cyclic AMP (cAMP), which integrates with progesterone signaling to regulate transcriptional activity. ^2^ Increases in cAMP are triggered by activation of Gαs-coupled G-protein coupled receptors (GPCRs) which promote adenylate cyclase activity to catalyze the conversion of adenosine triphosphate (ATP) to cAMP. However, identifying the upstream signaling molecules and receptors in the endometrium has proved problematic, and several factors, including relaxin, corticotrophin-releasing factor, human chorionic gonadotrophin (hCG) and prostaglandin E2 (PGE2) have been implicated. ^2^ Based on phylogenetic analysis, it is likely that PGE2 acting through its cognate GPCR EP2, encoded by *PTGER2*, is the main cAMP-inducing signal underlying decidualization. ^4^ Indeed, stimulation of endometrial stromal cells with PGE2 in combination with medroxyprogesterone (MPA, a progestin) is effective in eliciting decidual responses, as characterized by the induction and secretion of canonical markers, such as prolactin (encoded by *PRL*). ^4–6^ However, other factors, particularly relaxin, may also contribute. ^2,4,6–9^ Endometrial stromal cells are therefore exposed to a variety of hormones that induce cAMP signaling and yet still maintain the capacity to decode cAMP from specific receptors to selectively drive decidualization.

One accepted model that explains how cells translate multiple signals is via spatio-temporal mechanisms that specify and, in turn, enable diversification of downstream roles. ^10^ This includes the ability of Gαs-coupled GPCRs to generate cAMP signaling in distinct sub-cellular locations, creating localized cAMP pools that are differentially decoded. Thus, in addition to the archetypal activation of GPCRs and G-protein signaling at the plasma membrane, GPCRs can activate signaling following ligand-induced endocytosis to endosomes, as well as activation of G-proteins from the Golgi, mitochondria and nucleus. ^11,12^ There is further diversification of signals and functional responses within endosomes through differential targeting of GPCRs to early endosomes (EEs) or very early endosomes (VEE), compartments that are physically, biochemically and functionally distinct. ^13–15^ In terms of GPCR/G-protein signaling, trafficking of receptors such as the β2-adrenergic receptor to EEs contributes to the sustained, or second wave of Gαs/cAMP signaling, while other receptors such as the LHCGR, mediate their primary and acute Gαs/cAMP signaling response from the smaller VEEs. ^14^ This differential endosomal sorting is driven by the PDZ domain-containing protein GIPC (GAIP-interacting protein C-terminus), and in VEEs cAMP signaling is negatively regulated by the adaptor protein APPL1 (adaptor protein, phosphotyrosine interacting with PH domain and leucine zipper 1). ^14^ Overall, this has resulted in the model of ‘location bias’ for GPCR activity, whereby spatial localization of receptors directs differential downstream functions from the same G-protein/second messenger signal. ^16^

In this study, we sought to determine if candidate activators of cAMP signaling; PGE2 and relaxin, may be differentially regulated in primary human endometrial stromal cells (EnSCs). We identify that PGE2-mediated EP2 activation of cAMP signaling is less efficacious but more sustained compared to relaxin/RXFP1. While cAMP signals from both receptors require receptor internalization, PGE2/EP2 activity is regulated at a spatial level via the VEE/APPL1 compartment. We report that loss of the VEE machinery dysregulates PGE2-mediated cAMP production and crucial decidual signaling pathways, resulting in a disordered differentiation response. This underpins a pivotal role for endocytosis and spatial targeting of the EP2 receptor in coordinating the decidual gene network.

## Results

### PGE2 and relaxin induce distinct cAMP profiles

As depicted in figure 1A, we cultured primary EnSCs from whole endometrial biopsies to investigate how cells respond to PGE2 and relaxin to impact decidualization. Intracellular cAMP levels were quantified after 5 minutes exposure to increasing concentrations of PGE2 or relaxin, and dose-dependent responses were observed. LogEC_50_ values obtained were −6.70 ± 0.29 (200nM) and −7.29 ± 0.23 (51nM) for PGE2 and relaxin, respectively (mean ± SD), with maximal cAMP inductions achieved at ∼1µM (Figure 1B). Temporal cAMP profiles exhibited by both 1µM PGE2 and 1µM relaxin were sustained for up to 60 minutes and no additive effect was observed with co-addition (Figure 1C). However, at all time points, relaxin induced far greater levels of cAMP than PGE2. This difference in efficacy is unlikely to be due to differential regulation by phosphodiesterase enzymes, as both relaxin and PGE2-induced cAMP profiles were enhanced in the presence of the phosphodiesterase inhibitor, IBMX, yet their temporal profiles were relatively similar (Figure S1A).

**Figure 1.**
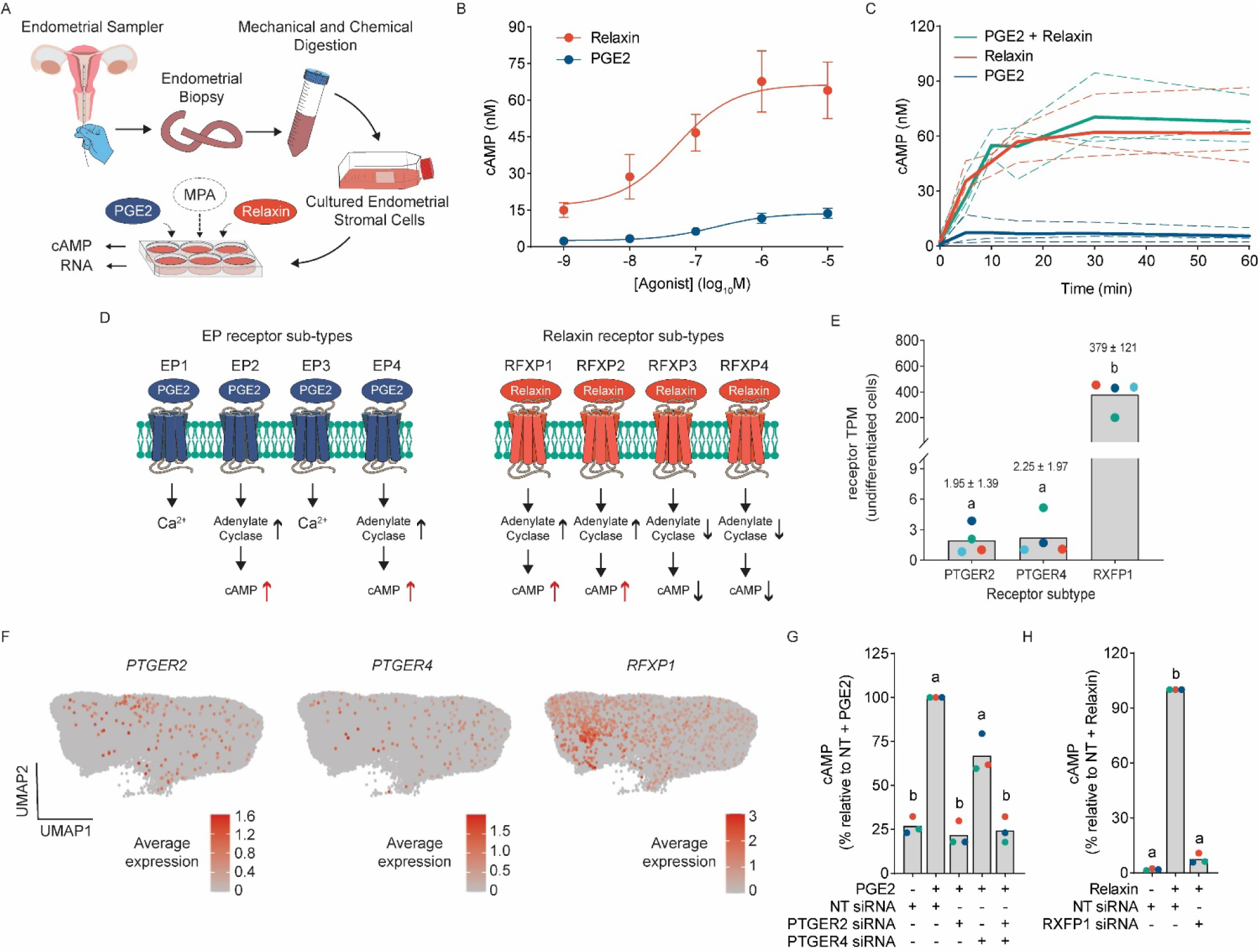
PGE2 and relaxin induce distinct cAMP profiles via EP2 and RXFP1. (A) Schematic depiction of experimental procedures to establish primary EnSC cultures from endometrial biopsies. (B) Concentration-dependent cAMP induction in EnSCs generated by a 5-minute stimulation with either PGE2 or relaxin. Data are mean ± SD, n=3. (C) Temporal cAMP profiles generated by either 1µM PGE2 or 1µM relaxin, or in combination. Plots from individual patients are represented by dashed lines with bold lines indicating mean values, n=3. (D) Schematic depiction of PGE2 (left panels) and relaxin (right panels) receptor subtypes and their associated signaling cascades. (E) Expression of *PTGER2*, *PTGER4* and *RXFP1* transcripts in EnSC cultures (GEO accession numbers GSE246591). Data points from individual patients are color-matched and shown together with mean TPM values ± SD. Different letters indicate statistical difference (*P*<0.05) between groups (ANOVA and Tukey’s multiple comparison test), n=4. (F) UMAPs depicting receptor expression in stromal cells from single-cell RNA sequencing of whole endometrial biopsies (GEO accession number GSE247962). Data are pooled from 12 individual patients. Positive cells and the level of receptor expression are shown by color key as indicated. (G) The relative change in cAMP to 5-minute stimulation with PGE2 in EnSCs following siRNA depletion of *PTGER2* and *PTGER4*. (H) The relative change in cAMP to 5-minute stimulation with relaxin in EnSCs following siRNA depletion of RXFP1. Data points from individual patients are color-matched and shown together with bar graphs denoting mean. Different letters indicate statistical difference (*P*<0.05) from NT siRNA, stimulated cells (ANOVA and Dunnett’s multiple comparison test), n=3.

As illustrated in figure 1D, PGE2 and relaxin can activate distinct GPCRs, with four receptor sub-types for both PGE2 (EP1-4) (encoded by *PTGER1-4*) and relaxin (RXFP1-4) (encoded by *RXFP1-4*). Each receptor sub-type has a unique expression pattern, G-protein coupling and signaling characteristics. ^17,18^ Of the PGE2 receptors, only EP2 and EP4 couple to Gαs to increase intracellular cAMP, and for relaxin this is limited to RXFP1 and 2. In primary EnSCs (GEO dataset GSE246591), *PTGER2*, *PTGER4* and *RXFP1* are expressed, but not *RXFP2* (Figure 1E). Of note, transcripts for *RXFP1* were 195- and 169-fold greater than *PTGER2* and *PTGER4*, respectively, which may in part underlie the higher levels of ligand-induced cAMP. Similarly, by using single-cell RNA sequencing data obtained from whole endometrial biopsies (GEO dataset GSE247962) we identified considerable cell heterogeneity in receptor expression in the stromal compartment, with only a small proportion of cells expressing *PTGER2* (0.57 %) and *PTGER4* (0.42 %), compared to a higher proportion of stromal cells expressing transcripts for *RXFP1* (3.12 %) (Figure 1F). RXFP2 was not detected. siRNA-mediated depletion of *PTGER2* or *PTGER4* demonstrated that EP2 was the primary receptor mediating PGE2-dependent cAMP in our cultures with minor contribution from EP4 (Figures 1G and S1B). Similarly, knockdown of *RXFP1* abolished relaxin-mediated cAMP signaling implicating this receptor subtype in the response (Figures 1H and S1C).

### PGE2, not relaxin, drives decidualization

Cultured EnSCs readily differentiate in response to cAMP and progestin, a protocol widely used to study decidualization. ^2^ We therefore treated EnSCs with either 1µM PGE2 or 1µM relaxin in combination with 1µM medroxyprogesterone acetate (MPA) (a progestin) for 4 days and measured the induction of decidual genes by RTqPCR. *IGFBP1* and *PRL* are recognized as canonical markers of decidualization, whereas the molecular chaperone protein clusterin, coded by *CLU,* and the IL-33 receptor, coded by *IL1RL1*, mark distinct cellular subtypes that emerge during differentiation. *CLU* expression predominates in progesterone-resistant cells, whereas *IL1RL1* marks progesterone-dependent decidual cells. ^19^ An increase in all decidual markers was observed following 4-day treatment with PGE2/MPA. However, and despite the robust and more efficacious cAMP signal induced by relaxin, minimal (*PRL*, *CLU*) or no (*IGFBP1*, *IL1RL1*) increases in decidual markers were observed with relaxin/MPA stimulation (Figure 2A). The data not only substantiates the role of PGE2 in inducing decidualization, but also highlights how increasing receptor-mediated cAMP levels, per se, is insufficient for stromal cell differentiation, emphasizing the importance of regulatory mechanisms that govern receptor signaling.

**Figure 2.**
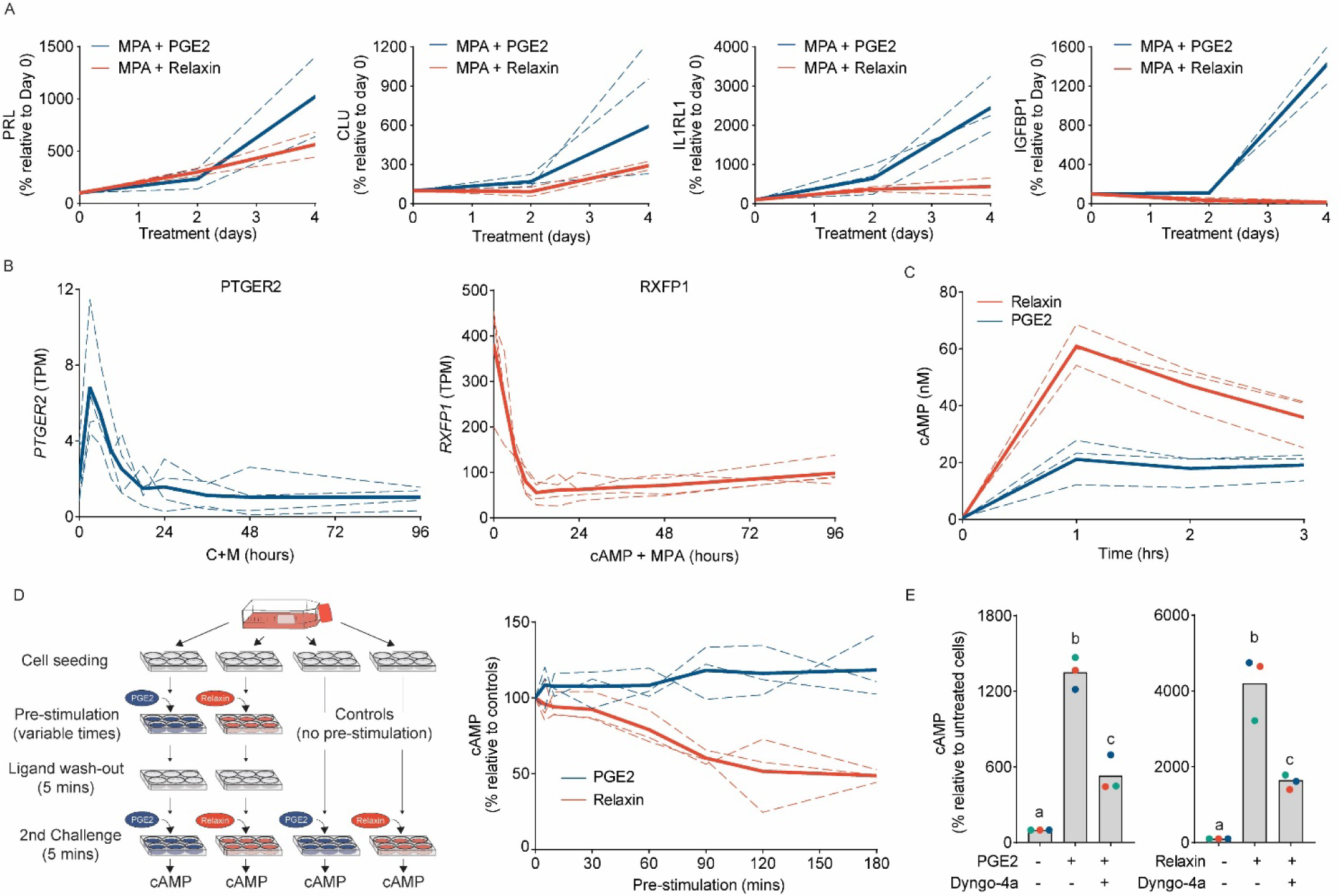
PGE2, and not relaxin, drives decidualization of EnSCs. (A) RTqPCR analysis of the relative changes in transcripts for decidual genes following 2 and 4-day treatment with PGE2 or relaxin in combination with MPA. (B) Changes in levels of transcripts for *PTGER2* (left panel) and *RXFP1* (right panel) in EnSCs treated with 8-bromo-cAMP and MPA from 3 hours up to 4 days (GEO accession number GSE246591), n=4. (C) Extended temporal cAMP profiles in EnSCs stimulated with either PGE2 or relaxin for up to 3 hours, n=3. (D) Schematic representation of the experimental procedures used for the resensitization assay (left panel). The magnitude of secondary cAMP induction in EnSCs following a pre-stimulation, desensitizing challenges with PGE2 or relaxin for variable times (right panel). Data are normalized as a percentage of control responses (no pre-stimulation). Plots from individual patients are normalized to the magnitude of cAMP signal in control (no pre-stimulation) cells, n=3. For A-D, individual cultures are represented by dashed lines with bold lines indicating mean values. (E) Induction of cAMP by PGE2 (left panel) or relaxin (right panel) following inhibition of receptor internalization with the dynamin-inhibitor, Dyngo-4A. Data points from individual patients are color-matched and shown together with bar graphs denoting mean. Different letters indicate statistical difference (*P*<0.05) from untreated, unstimulated cells (ANOVA and Dunnett’s multiple comparison test), n=3.

To examine differential regulation between EP2 and RXFP1 we analyzed RNA-seq data from EnSCs that had been decidualized with 8-bromo-cAMP (a receptor-independent cAMP analog) and MPA over an acute time-course from 3-96 hours (GEO accession number GSE246591), thus allowing us to temporally profile transcripts during the very early stages of decidualization. After an initial increase, levels of transcripts for *PTGER2* remained relatively stable up to 96 hours of treatment (Figure 2B, left panel). However, the number of transcripts for *RXFP1* dropped rapidly after initiation of treatment and were ∼85% less than undifferentiated cells by 12 hours (Figure 2B, right panel). With concern that this experiment used 8-bromo-cAMP instead of physiologically relevant inductions of cAMP, we characterized this regulation further by treating cells with 8-bromo-cAMP, relaxin or PGE2 alone, or in combination with MPA, for up to 4 days and quantified receptor expression via RTqPCR. For all treatments, expression of *PTGER2* was relatively stable, however *RXFP1* expression decreased markedly (∼80%) by day 2, but only when in the presence of MPA (Figure S2A). To ascertain if the cAMP signals downstream of the receptors are differentially regulated, we first measured cAMP levels in the continued presence of either ligand for up to 3 hours. The relaxin-mediated cAMP signal declined after 1 hour, suggesting a loss of signaling as receptors desensitize, but this was not apparent with PGE2 where signaling was sustained (Figure 2C). To further examine receptor desensitization, we assessed the ability of each ligand to induce signal resensitization. As depicted in Figure 2D, left panel, cells were first challenged with saturating concentrations of either 1µM PGE2 or 1µM relaxin for various timepoints and then washed for 5 minutes. Importantly, this wash was sufficient to return induced cAMP to basal levels (Figure. S2B). cAMP levels were then measured from a second acute (5 minutes) restimulation. Concordant with the sustained signal, the distinct temporal profiles indicated that PGE2/EP2-induced cAMP did not desensitize and could re-initiate cAMP signals following ligand washout, even after 3 hours ligand stimulation, while the relaxin-induced cAMP signal desensitized rapidly (Figure 2D, right panel). These observations highlight differential regulation between EP2 and RXFP1 receptors and offer plausible explanations for the poor induction of decidual genes with relaxin/MPA after 4 days.

One possible mechanism underpinning the sustained PGE2 signal profile is the ability of GPCRs to induce cAMP from endosomal compartments in addition to the plasma membrane. To assess the role of receptor internalization, cells were treated with dyngo-4A, a selective and potent dynamin GTPase inhibitor that blocks dynamin-dependent endocytosis. ^20^ Dyngo-4A blocked the internalization of agonist-stimulated, FLAG-tagged EP2 receptors, as visualized by confocal microscopy (Figure S2C). The cAMP signals from both PGE2 and relaxin were partially, but significantly, inhibited by dyngo-4A pre-treatment confirming an important functional role for receptor internalization and endosomal signaling (Figure 2E).

### EP2 receptors are internalized through very early endosomes

We have demonstrated previously that certain GPCRs are targeted to VEEs, which are biochemically, physically and functionally distinct from EEs. ^13,14^ As depicted in figure 3A, the VEE compartments are smaller than EEs and devoid of classical endosomal markers such as Rab5, EEA1 and PI3P. Some GPCRs are directed to these compartments via GIPC to activate signaling and be rapidly recycled, whilst APPL1 localizes to a subpopulation of VEEs to negatively regulate endosomal GPCR/G-protein signaling and sort receptors for rapid recycling. ^14^ Collectively this leads to a sustained pattern of G protein/second messenger signaling. We postulated that the sustained cAMP signal from PGE2/EP2 was driven by trafficking through the VEEs. To study endosomal targeting, FLAG-tagged EP2 receptors (FLAG-EP2) were transfected into EnSCs and visualized via confocal microscopy. Trafficking was directly compared to FLAG-tagged β2-adrenergic receptors (FLAG-β2AR), a receptor known to internalize to EEs. ^13,21,22^ Both receptors internalized within 5 minutes of agonist stimulation and appeared within endosomes. However, there was a clear difference in the size of their respective endosomal compartments (Figure 3B). Quantification of endosome size revealed that FLAG-β2AR enters compartments 0.70 nm ± 0.23 in diameter, whereas compartments containing FLAG-EP2 were 0.49 nm ± 0.17 (data are mean ± SD, *P*<0.0001 via Mann-Whitney test) (Figure 3B, right panel). In addition, a subpopulation of internalized EP2 colocalized with APPL1 positive compartments (Figure 3C). These characteristics are consistent with observations describing VEEs and confirm internalization of EP2 into these compartments. ^13,14^

**Figure 3.**
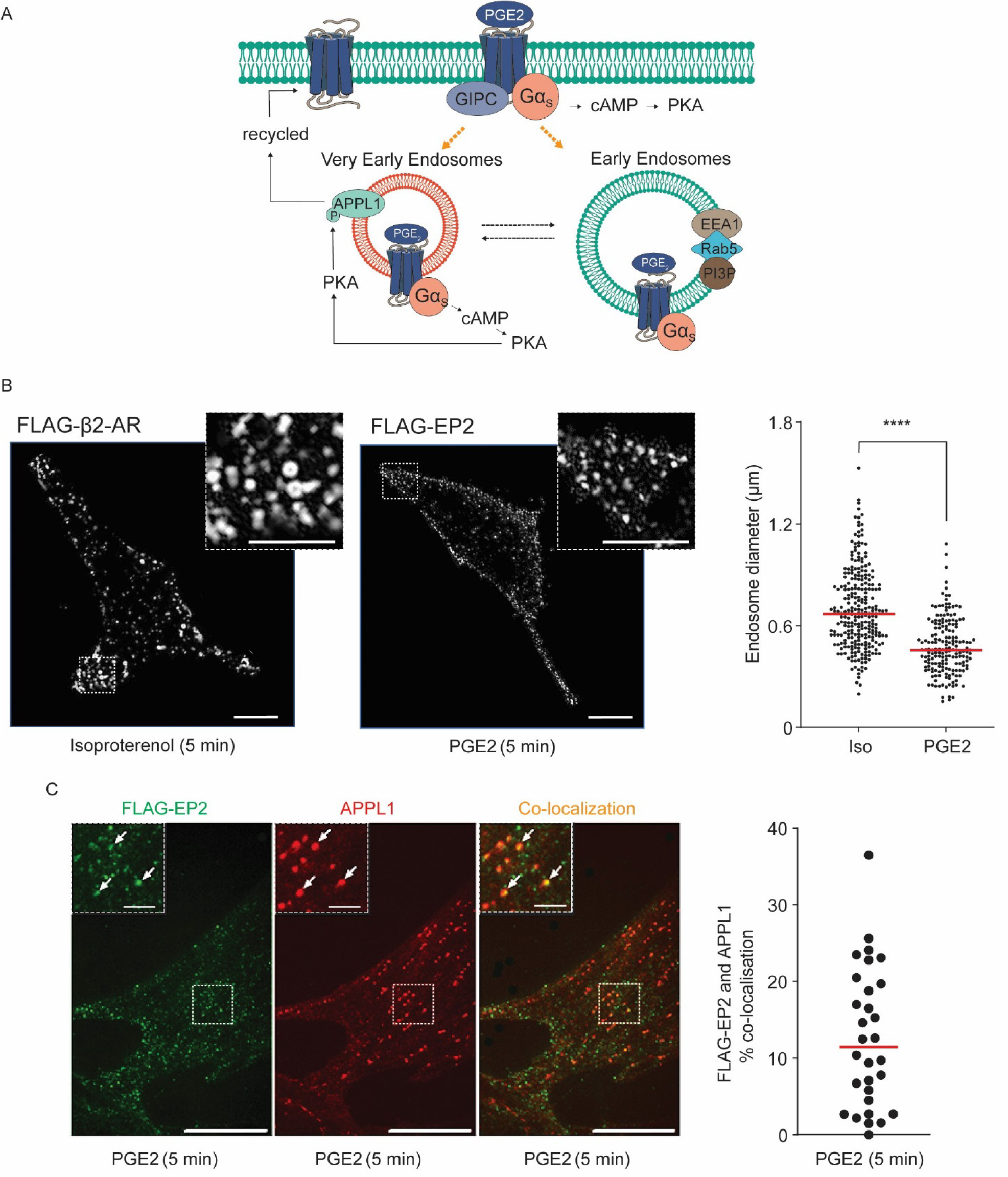
Internalized EP2 receptors traffic through very early endosomes. (A) Schematic representation of endosomal compartments and their cargo. (B) Representative images from FLAG-βAR and FLAG-EP2 as captured by TIRF microscopy, 5 minutes post isoproterenol and PGE2 stimulation, respectively. Quantitation of endosome diameter (right panel). 182 endosomes (FLAG-EP2) and 264 endosomes (FLAG-βAR) were measured from 4 independent cultures. **** denotes a *P*-value of <0.0001 via Student’s t-test. (C) FLAG-EP2 (green) and APPL1 (red) positive endosomes with co-localization detected as orange staining, 5 minutes post PGE2 stimulation (left panels) and quantitation of overlay against the average of all cells (right panel). Arrows indicate FLAG-EP2 and APPL1 positive endosomes. Data are obtained from 30 representative regions chosen at random, n=3. Scale bars = 20µm, inset = 10µm.

### PGE2, but not relaxin, -mediated cAMP signaling is regulated by APPL1 and GIPC

To determine if cAMP induction by PGE2/EP2 were regulated by VEE machinery, levels of GIPC or APPL1 were depleted from EnSCs by siRNA (Figure 4A). We have previously demonstrated that GIPC knockdown redirects receptors destined for VEEs into the EEs, whereas APPL1 knockdown enhances signaling and also prevents receptor recycling, leading to retention of receptors within the VEEs. ^13,14,23^ Consistent with these known roles, the acute induction of cAMP in response to PGE2, but not to relaxin, in APPL1-depleted cells increased ∼2 fold (Figure 4B). Depletion of GIPC had no significant effects on responses to either ligand. Both APPL1 and GIPC are known to impact receptor recycling from VEEs via distinct mechanisms, but these ultimately lead to a post-endocytic sorting pathway that facilitates receptor resensitization. ^13,14^ To assess if depletion of GIPC or APPL1 impacts the ability of EP2 to resensitize and thus promote cAMP signal recovery, we assessed cAMP induction to repeated PGE2 challenge (Figure 4C, left panel). Knockdown of either GIPC or APPL1 inhibited the ability of EnSCs to respond to a second PGE2 challenge, indicating that both are important for signal resensitization (Figure 4C, right panel). Overall, this data demonstrates that PGE2 signaling by EP2 employs VEE-associated machinery to fully re-activate and maintain sustained cAMP signaling.

**Figure 4.**
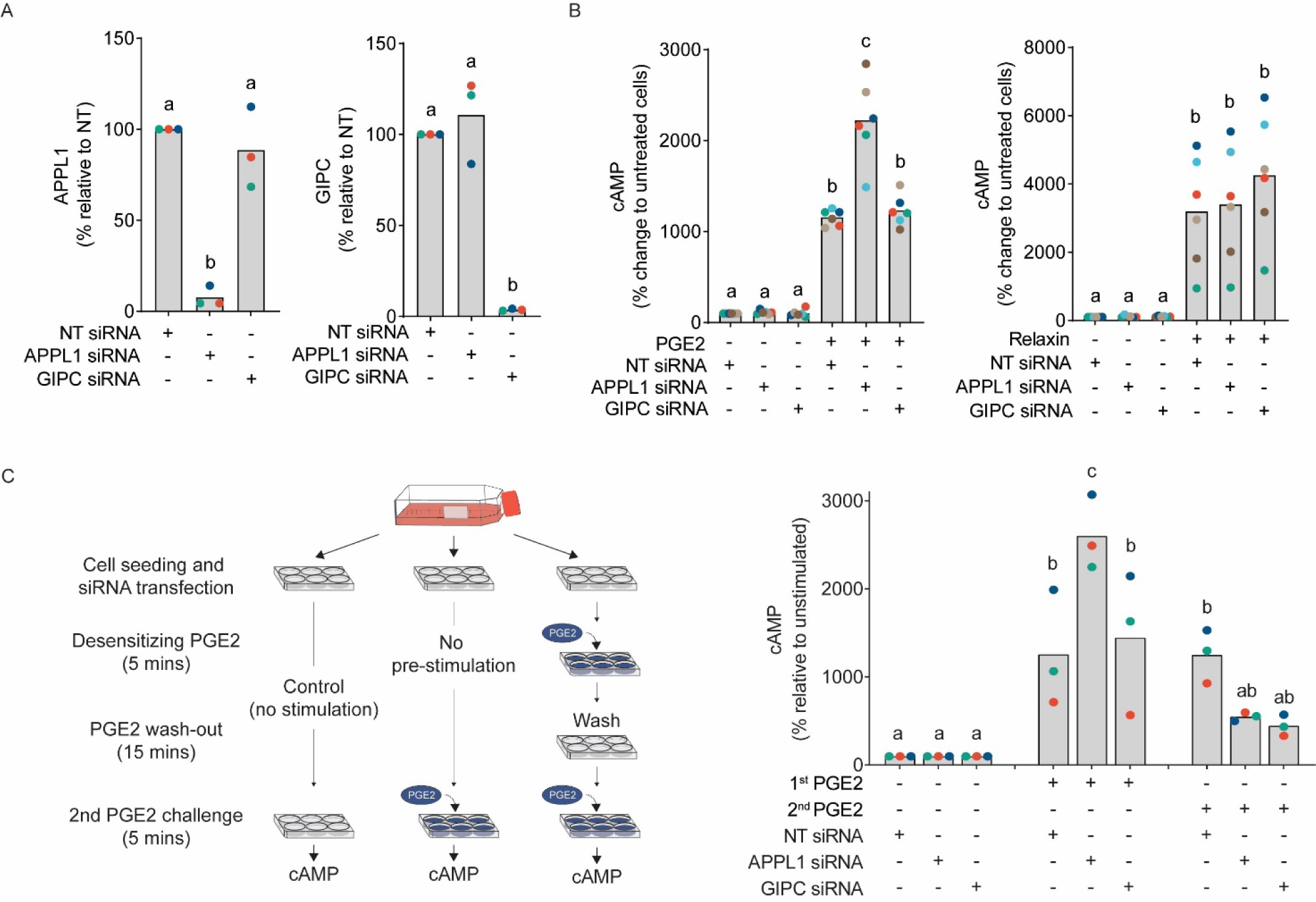
PGE2, but not relaxin, mediated cAMP responses are regulated by APPL1 and GIPC. (A) RTqPCR analysis of transcripts for APPL1 (left panel) and GIPC (right panel) following their depletion in EnSCs by siRNA. Data from individual patients are color-matched and shown with bar graphs denoting mean values. Differing letters indicate significance from NT siRNA controls (*P*<0.05) (ANOVA and Dunnett’s multiple comparison test, n=3). (B) Induction of cAMP after 5-minute stimulation with PGE2 (left panel) and relaxin (right panel) following depletion of APPL1 and GIPC by siRNA. Data from individual patients are color-matched and shown with bar graphs denoting mean values. Differing letters indicate significance from NT siRNA, unstimulated controls (*P*<0.05) (ANOVA and Dunnett’s multiple comparison test, n=6). (C) Schematic representation of experimental procedures used to assess resensitization of EP2 receptors depleted of APPL1 and GIPC (left panel). The cAMP signal from a 5-minute PGE2 challenge (2^nd^ response) following an identical desensitization challenge (1^st^ response) or control, and ligand washout (right panel). Data from individual patients are color-matched and shown with bar graphs denoting mean values. Differing letters indicate significance from NT siRNA, unstimulated controls (*P*<0.05) (ANOVA and Dunnett’s multiple comparison test, n=3).

### Loss of APPL1 and GIPC inhibits decidualization

Given APPL1 and GIPC are required to maintain PGE2-induced cAMP, and that PGE2, and not relaxin, is a driver of stromal cell decidualization, we assessed the impact of APPL1 and GIPC knockdown on PGE2-mediated decidualization. To investigate this, three independent EnSC cultures were depleted of APPL1 or GIPC and treated with 1µM PGE2 and MPA for 4 days before total RNA was extracted and sequenced (Figure 5A). There was no loss of EnSC viability after 4-day treatment, as assessed by XTT assay (Figure 5B), and APPL1 and GIPC expression remained depleted for the duration of the experiment (Figure 5C). Of note is the significant increase in transcripts for GIPC with APPL1 siRNA (Figure 5C, right panel), suggesting a compensatory or corrective response, or indeed that GIPC is part of the gene network regulated by APPL1.

**Figure 5.**
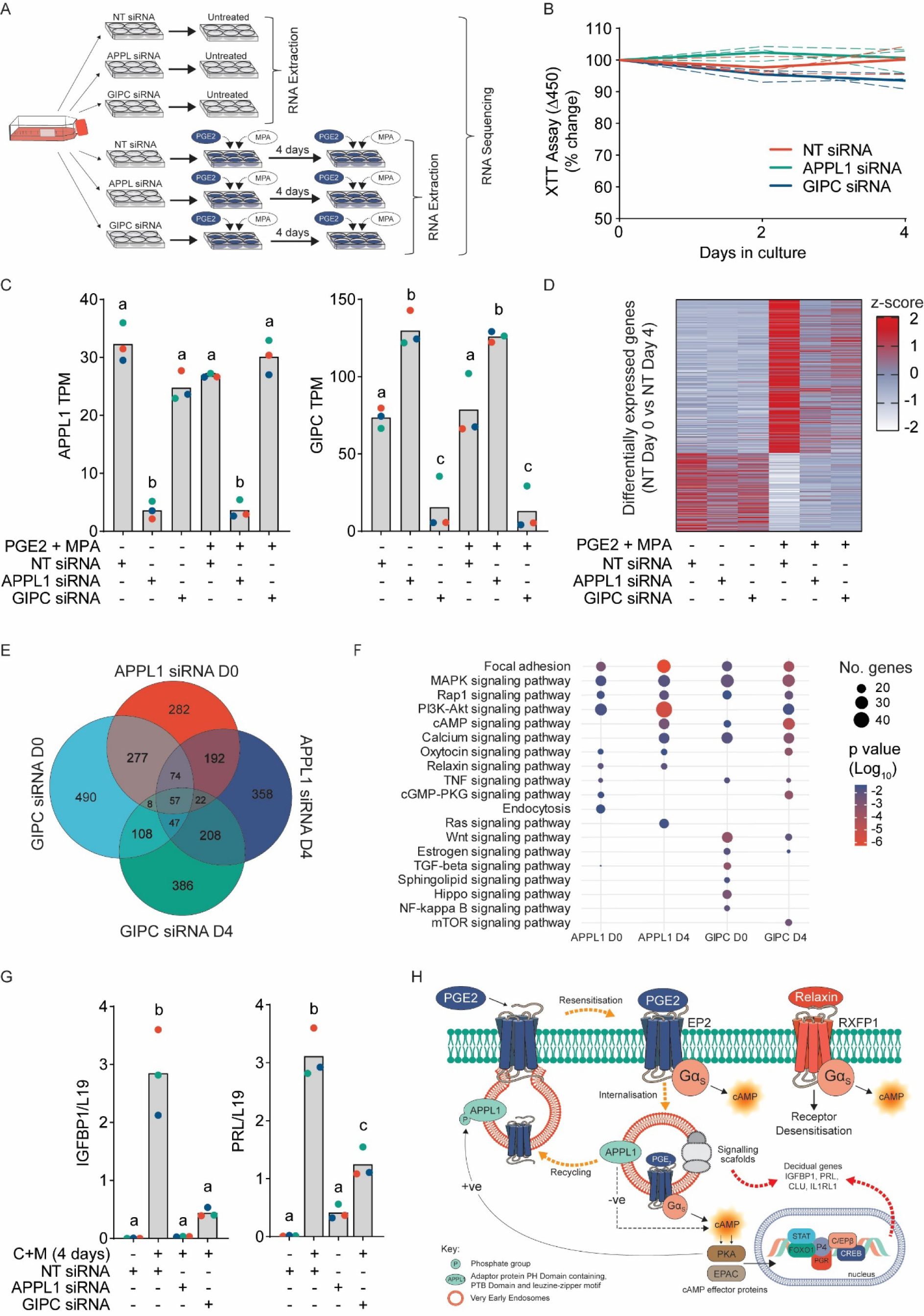
Loss of APPL1 and GIPC inhibits decidualization. (A) Schematic depiction of treatment protocol for RNA sequencing in EnSCs. (B) Relative changes in absorbance from XTT assay in untreated EnSCs depleted of APPL1 and GIPC for 4 days. Individual patients are shown as dashed lines, with mean values depicted by bold lines, n=3. (C) RTqPCR analysis of APPL1 (left panel) and GIPC (right panel) 4 days following siRNA depletion. Data from individual patients are color-matched with bar-graphs denoted mean values. Differing letters indicate significant between groups (*P*<0.05) (ANOVA and Tukey’s multiple comparison test, n=3). (D) Heat map of differentially expressed genes (Bonferroni correction, *P*<0.05) identified from bulk RNA-sequencing between untreated NT siRNA cells and those treated with PGE2/MPA for 4 days. A total of 645 differentially expressed genes were identified (435 upregulated and 210 downregulated) and each gene is scaled (Z-scored) across treatments to show changes with APPL1 and GIPC depletion. Red, blue and white colors represent high, medium and low expression, respectively, as per key. Data are n=3. (E) Venn diagram depicting the number of individual and common differentially expressed genes (Bonferroni correction, *P*<0.05) identified comparing APPL1 and GIPC depleted EnSCs to treatment-matched NT siRNA cells. (F) Selected Kyoto Encyclopedia of Genes and Genomes (KEGG) pathway enrichment of differentially expressed genes (full lists of KEGG enrichment terms are available in Tables S2-5). The size of circles is relative to the number of genes in each enrichment term, and the color represents *P*-value calculated as a result of enrichment degree. (G) RTqPCR analysis showing relative changes in decidual genes *IGFBP1* (left panel) and *PRL* (right panel) following 4-day treatment with 8-bromo-cAMP and MPA (C+M) in EnSCs depleted of APPL1 or GIPC. Data from individual patients are color-matched with bar-graphs denoted mean values. Differing letters indicate significant compared to untreated (*P*<0.05) (ANOVA and Dunnett’s multiple comparison test), n=3. (H) Summary schematic detailing the rapid recycling and resensitization of EP2 receptors through the VEEs and its contribution to decidualization of EnSCs.

We identified a total of 645 differentially expressed genes (Bonferroni adjusted *P*-value <0.05) in EnSCs between non-targeting (NT) D0 and NT D4 PGE2/MPA-treated cells, of which 435 (67%) were upregulated and 210 (33%) downregulated (Figure 5D). The most highly upregulated gene was *MAOB* (adjusted *P*-value = 2.16 x 10^−140^), encoding monoamide oxidase B, a metabolic enzyme with varied roles in metabolism and proliferation, and reported roles in embryo receptivity. ^24,25^ Many other genes that form part of the decidual gene network were upregulated suggesting a strong decidual response. Select examples include *PRL* (9.2 x 10^−27^), *FOXO1* (4.13 x 10^−17^), *IL1RL1* (7.87 x 10^−16^), HAND2 (1.65 x 10^−20^), *SCARA5* (2.43 x 10^−11^), ZBTB16 (1.11 x 10^−10^), *CXCL14* (2.58 x 10^−8^) KLF4 (1.66 x 10^−8^) and *CDKN1C* (2.98 x10^−4^) (Figure S3A). The induction of *IGFBP1* transcripts was evident but non-significant. Strikingly, knockdown of either APPL1 or GIPC abolished the induction of most upregulated genes, and at least partially recovered those that were downregulated. Out of the 435 genes upregulated with PGE2/MPA treatment, 185 were significantly downregulated in parallel cells depleted of APPL1, and 154 with GIPC (Bonferroni adjusted *P*<0.05), indicating a wide and universal blunting of the decidual response (Figure 5D).

Even when applying the conservative Bonferroni method to correct for multiple comparisons, we identified a large number of differentially expressed genes with APPL1 and GIPC depletion across both untreated (D0) and treated (D4) cells (Figure 5E), the majority of which were downregulated (Figure S3B). In all cases, comparisons were made against treatment-matched NT siRNA transfected cells. Of note, is the large number of shared differentially expressed genes between APPL1 and GIPC depleted cells and between D0 and D4 cells, suggesting perturbation of common pathways (Figure 5E). Pathway analysis of differentially expressed genes was performed using the Kyoto Encyclopedia of Genes and Genomes (KEGG) database and revealed dysregulation of several important signaling pathways, including cAMP and Rap1, a down-stream regulator of EPAC1/2. Like PKA, EPAC proteins directly interact with cAMP as effector proteins, translating cAMP signals to diverse biological functions. Other conspicuous GO categories such as MAPK and PI3K-Akt signaling were also common across treatment groups, and many others such as TNF and TGFβ, and Wnt signaling were also significant. All these pathways regulate diverse cellular functions such as gene expression, differentiation and cell fate, and all are pivotal to decidualization. ^2,26^ In addition, focal adhesion was a highly enriched KEGG term across all treatments. Focal adhesions form an integrin-rich bridge between the cytoskeleton and the extracellular matrix and co-ordinate responses to tensile stresses and extracellular signals. They are critically dependent on endosomal transportation for protein turnover and spatial organization of integrins. ^27–29^ They have also been implicated in pregnancy recognition and implantation. ^30–32^ A full list of KEGG terms for each comparison is available in tables S2-5. As well as shared transcriptomic changes, there were differentially expressed genes unique to cells depleted of either APPL1 or GIPC (Figure S3C), and we posited that this reflected distinct and divergent functions for different endosomal compartments. Indeed, at day 4, select KEGG pathways associated with negatively regulated genes in APPL1 depleted cells included MAPK, mTOR and Rap1 signaling, whereas those for GIPC included relaxin signaling and the PI3K-Akt pathway (Figure S3D). In fact, the effect on PI3K-Akt signaling was reciprocal as this was associated with upregulated genes after APPL1 knockdown.

The dysregulation of notable decidual pathways in unstimulated cells indicates basal turnover or receptor trafficking through paracrine responses. We therefore postulated that dysregulation of the endosomal pathways whilst treating EnSCs with 8-bromo-cAMP (a deciduogenic cue that by-passes receptors and endosomes) and MPA would also moderate the decidual response. Indeed, in this experiment, APPL1, and to a lesser degree GIPC, depletion abolished the induction of *IGFBP1* and *PRL* (Figure 5E). As illustrated in figure 5G, we therefore propose that signaling through the VEEs is essential for decidualization, firstly by rapidly resensitizing cell responsiveness and sustaining intracellular cAMP, and secondly by spatially organizing effector proteins to co-ordinate key decidual signaling pathways.

## Discussion

Decidualization requires convergence of progesterone and cAMP signaling, and despite many purported signaling molecules, PGE2 and relaxin remain likely candidates as the source of cAMP. They exert their effects by activating respective EP2 and RXFP1 Gαs-coupled GPCRs, however little is known about how endometrial stromal cells decode and interpret these signals to elicit appropriate responses. The significance of sub-cellular location in receptor signaling has emerged as a crucial model for both diversifying and specifying G-protein signaling pathways from various GPCRs. In this study, we identified the distinct spatial regulation receptors of EP2 and RXFP1 receptors and assessed how differential regulation of cAMP impacts decidualization in endometrial stromal cells.

PGE2 is a ubiquitous paracrine signaling molecule with diverse roles in many systems. It is derived from arachidonic acid in a multi-step process through Phospholipase A2 and COX-2 (PTGS2) enzymes. COX-2/PTGS2 expression is common among all endometrial cell types with high levels in epithelial cells. ^33–36^ Relaxin is principally released from the corpus luteum, a temporary structure formed in the ovary after ovulation, but is also synthesized locally within the endometrium. ^37^ Both signals are therefore present at time of decidualization. Despite relaxin inducing a more efficacious cAMP signal than PGE2 in EnSCs, we found that PGE2, and not relaxin, promoted decidualization when cells were co-treated with a progestin. We therefore investigated whether receptor endocytosis and sub-cellular location in receptor-mediated cAMP signaling, rather than the magnitude of induction, was an important mechanism underlying decidualization.

Intracellular compartmentalization of signaling provides a system for common upstream second messenger signals, such as cAMP, to orchestrate diverse roles in the same cell. ^11,12^ In this study, we propose that relaxin and PGE2 signals are differentially internalized and spatially regulated. The cAMP induced by both ligands were partially sensitive to inhibition of receptor internalization, suggesting both can signal from the plasma membrane and endosomal compartments. Using FLAG-tagged human EP2 receptors to monitor receptor trafficking we found that PGE2 was able to induce EP2 internalization in EnSCs. This contrasts with previous reports in HEK 293 cells where no PGE2-induced endocytosis of EP2 was observed. ^38^ There may, therefore, be distinct machinery driving receptor internalization in endometrial stromal cells that provides a means for these cells to decode PGE2-mediated signals and mount a precise and targeted response. Indeed, differences in levels of GPCR kinases have been proposed to underlie differential internalization of other GPCRs such as the μ-opioid receptor across different cell types. ^39^ EP2 internalized into small endosomes that were partially positive for APPL1 and generated cAMP signals that were regulated by GIPC and APPL1; all features of a receptor that is internalized and regulated through very early endosomes (VEEs). Conversely, relaxin-induced cAMP was unaffected by the loss of the GIPC and APPL1, indicating sorting to distinct compartments. The PDZ protein GIPC was important in maintaining the PGE2 cAMP signal via resensitization of cells, consistent with its role in sorting GPCRs to the VEE to facilitate rapid recycling back to the plasma membrane. ^13^ For receptors such as the LHCGR and β1-adrenergic receptor, GIPC binds to residues within the distal portion of its C-terminal tail. ^13^ While no PDZ ligand has been identified in the EP2 C-tail, the PDZ ligand predictor interface (a structure-based predictor for PDZ domain-peptide interactions) suggests only the last 5 residues (KQADL) may constitute a PDZ protein binding domain. ^40^ However, APPL1 is known to interact with GIPC, ^41,42^ and it is also plausible that GIPC exerts its actions on EP2 via APPL1. Depletion of APPL1 augmented PGE2-mediated cAMP induction consistent with its role as a negative regulator of VEE G-protein signaling for other GPCRs, such as those for reproductive hormones (LHR, FSHR) and short chain fatty acid receptors (FFA2). ^14,23^

Our data demonstrates that PGE2/EP2 signaling is tightly controlled through precise spatially and temporally organized cAMP signaling, and that perturbation of endosomal trafficking, via knockdown of GIPC or APPL1 lead to disruption of decidual gene networks. KEGG pathway analysis of differentially expressed genes highlighted dysregulation of key decidual signaling pathways including MAPK and PI3K/Akt. These, along with other significant KEGG terms such as TNF and TGFβ, and Wnt signaling, are all indispensable for decidualization, ^2^ and highlights a critical link between VEE signaling and engagement of EP2 with these pathways. Endosomes are known for their scaffolding properties, physically orientating internalized receptors to form complex assemblies with signalling molecules, ^43–45^ thus disruption of endosomal trafficking, combined with the sensitive and saturable nature of endosomally-induced signals, would affect activation of down-stream substrates. Indeed, active Akt can only phosphorylate certain substrates when compartmentalized into specific intracellular locations. ^46^ Similar KEGG pathways were also identified in unstimulated APPL1 and GIPC depleted cells suggesting these adaptor proteins modulate decidual gene expression through either receptor-independent mechanisms, or through basal and autocrine activity. Decidualization was also impaired when PGE2/EP2 signaling was bypassed with 8-bromo-cAMP, a membrane permeable cAMP analog, further emphasizing how large, sustained levels of cAMP alone are insufficient for decidualization.

Overall, we have identified an important mechanism in endometrial biology that explains how spatial regulation of EP2 generates cAMP to drive decidualization. Given the key role of endometrial decidualization in both establishing early pregnancy and placentation, dissecting the molecular mechanisms of PGE2-mediated decidualization will likely pave the way for improved understanding of its pathophysiological actions.

## Limitations

There is currently a lack of sensitive antibodies against EP2 to image endogenous receptor, thus internalization was monitored using an expressed FLAG-tagged EP2 receptor. We also have no direct evidence of native EP2 G-protein signaling from endosomes and rely on manipulation of endocytic pathways through depletion of trafficking proteins and the analysis of down-stream signals. A more specific, yet challenging approach would be to manipulate receptors through site-directed mutagenesis to disrupt signaling and protein interactions. At present, however, we don’t know which residues dictate VEE targeting.

## Acknowledgments

The authors are forever indebted to the women who volunteer and donate samples for research as well as the clinical staff that facilitate their involvement.

## Author contributions

Conceptualization, P.J.B., A.H and J.J.B; Methodology, P.J.B., A.W, A.H, J.J.B; Investigation, P.J.B., A.W, O.M; Data curation, P.J.B, A,W, O.M, C-S.K, E.L, P.V, Formal analysis, P.J.B, C-S.K, P.V, E.L; Writing – original draft, P.J.B., A.H., A.W and J.J.B; Writing - review and editing P.J.B., A.W., O.M, E.L, P.V, A.H, J.J.B. Funding acquisition, A.H and J.J.B; Resources, J.J.B; Supervision, P.J.B., A.H and J.J.B.

## Data and materials availability

Data associated with this study are present in the paper. In addition, expression data from RNA sequencing have been submitted to the Gene Expression Omnibus (GEO) repository (accession number: GSE246497).

## Competing interests

The authors have no competing interests to declare.

## Funding

This work was supported by funds from the BBSRC to J.J.B (BB/S00193X/1) and A.H. (BB/S001565/1).

## Experimental Procedures

### Key resource table

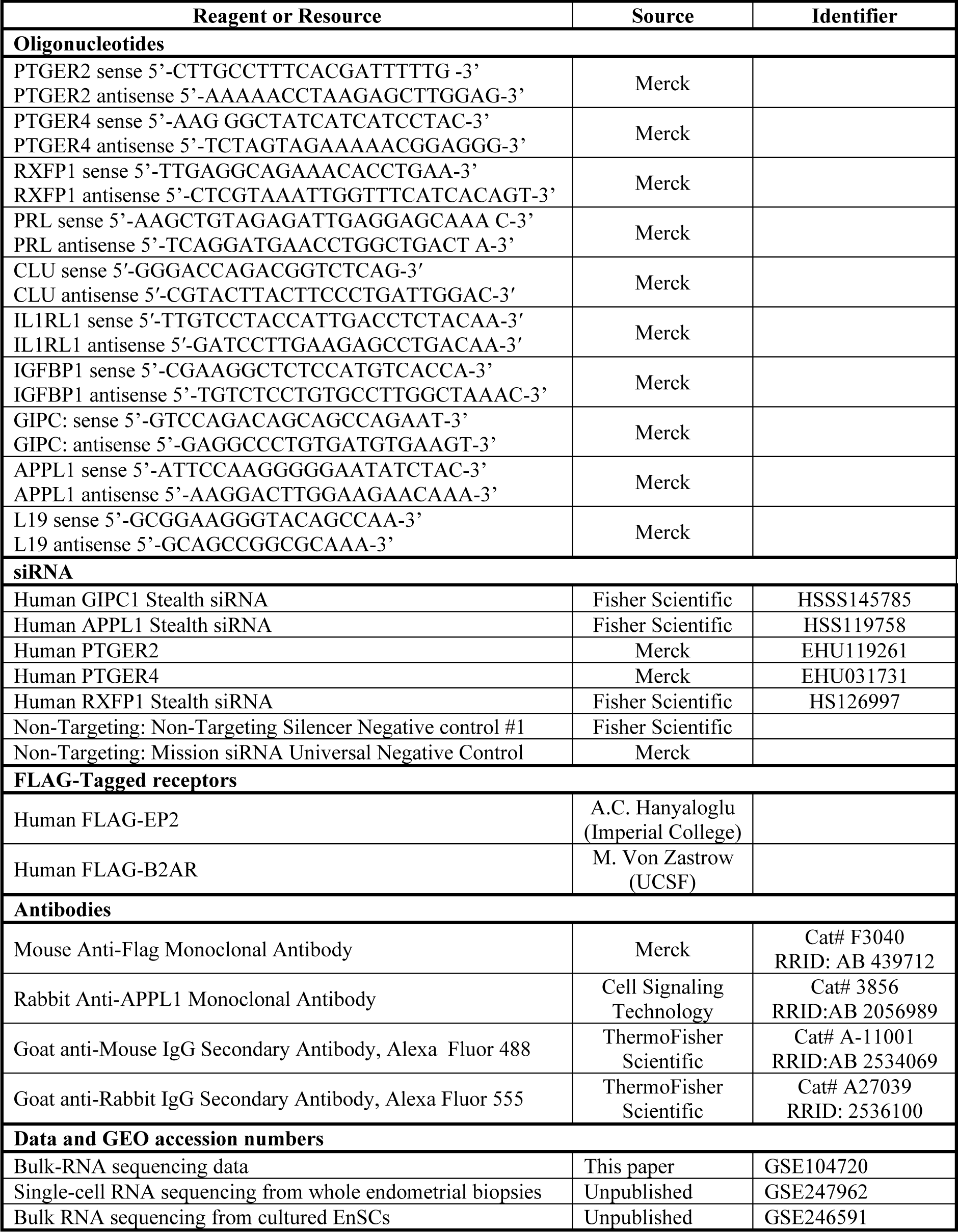

### Resource availability

#### Lead contact

Further information and reasonable requests for reagents should be directed to the lead contact, Dr. Paul Brighton. FLAG-EP2 is available from Professor Aylin Hanyaloglu under a materials transfer agreement with Imperial College London. This study did not generate new reagents.

#### Data and code availability

The RNA-seq data have been deposited in GEO and this paper analyses publicly available data sets. GEO accession numbers are listed in the key resources table. Any additional information required to reanalyze the data reported in this study is available from the lead contact upon request.

### Collection of human endometrial samples

Endometrial biopsies were collected for research purposes under the approval of the NHS National Research Ethics – Hammersmith and Queen Charlotte’s and Chelsea Research Ethics Committee (REC reference: 1997/5065) and Tommy’s National Reproductive Health Biobank (REC reference: 18/WA/0356), with written informed consent obtained from patients prior to tissue collection in accordance with the guidelines of the Declaration of Helsinki, 2000. Endometrial biopsies were obtained using a Endosampler (Medgyn, IL, U.S.A.) from patients attending a dedicated research clinic at the University Hospitals Coventry and Warwickshire (UHCW) National Health Service (NHS) Trust, Coventry, UK. Samples were timed to the secretory phase of the cycle as reported by patients using a home urinary LH test to date the pre-ovulatory LH surge. Appointments were timed 5–10 days later (LH+ 5-10). In total, 48 biopsies were used in the investigation to establish EnSC cultures. Demographic details of patients who donated biopsies are detailed in table S1.

### Endometrial tissue dissociation and stromal cell culture

#### EnSC dissociation and culture

Endometrial biopsies were transferred into additive-free DMEM/F12 media and processed immediately. Tissue was mechanically diced into a pulp with a surgical scalpel, and EnSCs were dissociated from endometrial biopsies by enzymatic digestion of the extracellular matrix using 500 µg/mL collagenase type Ia (Merck Life Sciences UK Ltd, Gillingham, UK) and 100 µg/mL DNase I (Lorne Laboratories Ltd, Reading, UK) diluted in additive-free DMEM/F12 media, for 1 hour at 37°C with manual agitation every 20 minutes. Digested tissue was flushed through a 40µm cell strainer to remove glandular cell clumps and undigested material, and the flow-through collected and cultured in DMEM/F12 supplemented with 10% dextran-coated charcoal-treated fetal bovine serum (DCC-FBS), antibiotic-antimycotic mix (100 units/mL of penicillin, 100 µg/mL of streptomycin, 250 ng/mL of Amphotericin B), 10µM L-glutamine (all Thermo Scientific, Loughborough, UK), 1 nM β-estradiol and 2 μg/mL insulin (Merck). Any red blood and unattached cells were removed by media change within 18 hours. Cells were cultured at 37°C in a 5% CO_2_, humidified environment and were lifted with 0.05% trypsin (5 minutes, 37°C), counted using a Neubauer-approved hemocytometer, and seeded as required. Experiments were performed at passage 2.

#### Decidualization

Confluent EnSC monolayers in 6-well plates were downregulated in phenol-free DMEM/F12 media containing 2% DCC-FBS, L-glutamine and antibiotic/antimycotics and decidualized with 1µM PGE2 or 1µM relaxin-2 (accession # P04090) (both Biotechne Ltd, Abbington, U.K.) in combination with 1µM medroxyprogesterone acetate (MPA) (Merck) for up to 4 days, with media refreshed every 2 days. Where required, cells were treated with 0.5 mM 8-bromo-cAMP (Merck), in combination with MPA. Further details for cell dissociation and culture can be found elsewhere. ^47^

#### siRNA Depletion

EnSCs were transfected using the jetPRIME transfection kit (Polyplus, Illkrich, France) exactly as per manufacturer’s instructions. 70% confluent EnSC monolayers in 6-well plates were downregulated overnight in 2% DCC-FBS/DMEM/F12 media and transfected with 50nM Stealth siRNA against human *GIPC1* or *APPL1* (Fisher Scientific, Loughborough, U.K.). For depletion of receptor expression, cells were transfected with 50nM Stealth siRNA against human *RXFP1* (Fisher Scientific), 50nM Mission esiRNA against human *PTGER2* or *PTGER4* (Merck). 50nM non-targeting Silencer™ Negative Control #1 siRNA (Fisher Scientific) or Mission siRNA Universal Negative Control (Merck) were used as controls. Further details are available in the Key Resource Table. Media was replaced 18 hours post-transfection and cells cultured for a further 2 days before experimental treatments. Knockdown was confirmed via RTqPCR.

### in silico analysis

Receptor expression data were derived from publicly available RNA sequencing data sets within the Gene Expression Omnibus (GEO) repository. Expression of transcripts for receptors in EnSC cultures was mined from bulk RNA sequencing GEO accession number GSE246591, and within the stromal compartment of whole endometrial biopsies from single-cell RNA sequencing data, GEO accession number GSE247962.

### Induction of cAMP

The induction of cAMP was quantified in EnSCs using a modified protocol of the High-Throughput Time Resolved Fluorescence (HTRF) cAMP Gαs dynamic kit (CisBio Bioassays, Codolet, France). A 2-step protocol allowed culture and stimulation of cells in 6-well plates and ensured cAMP levels were within the manufacturer’s detection range. Confluent EnSC cultures in 6-well plates were downregulated in 2% DCC-FBS media overnight, and then washed and allowed to equilibrate for 15 minutes in Krebs’–Henseleit Buffer (composition: NaCl: 118 mM, d-glucose: 11.7 mM, MgSO4·7H2O: 1.2 mM; KH2PO4 1.2 mM, KCl 4.7 mM, HEPES 10 mM, CaCl2·2H2O: 1.3 mM, pH 7.4, 37 °C). Cells were stimulated with either PGE2 or relaxin as per experimental requirements, and reactions terminated by rapid aspiration of buffer and immediate addition of 50 µl ice-cold lysis and detection buffer. 10 µl cell lysates were transferred to 384-well white walled, white bottomed plates and cAMP determined using the anti-cAMP cryptate donor and cAMP-d2 acceptor reagents exactly as per manufacturer’s instructions. Florescence was measured on a PHERAstar FS plate reader (BMG Labtech Ltd, Ortenberg, Germany). Where required, phosphodiesterase activity was inhibited by 300µM IBMX in Krebs’-Henseleit buffer, and dynamin by a 30 minutes, 30µM dyngo-4a (Abcam PLC, Cambridge, U.K.) pre-treatment prior to stimulation.

### RTqPCR

Total RNA was from extracted from cultured EnSCs and purified through columns using the RNeasy Plus Universal Mini kit exactly as per manufacturer’s instruction (Qiagen, Manchester, U.K.). Recovered RNA in nuclease-free water was analyzed on a Nanodrop spectrophotometer where concentration and purity (230/280 and 260/280 nm) were determined. Equal quantities of RNA were transcribed into cDNA using QuantiTect Reverse Transcription kits (Qiagen), and analysis of targets gene expression was performed on a 7500 Real-Time PCR System (Applied Biosystems, CA, U.S.A.) using Power SYBR Green PCR Master Mix (Life Technologies, Paisley, U.K.). The expression level of each gene was calculated using the ΔCT method and normalized against the expression of the L19 housekeeping gene. Primer sequences for all genes are listed in the key resources table.

### Microscopy

#### Receptor internalization and endosome size

EnSCs were cultured on glass coverslips prior to transient transfection with FLAG-EP2 or FLAG-β2AR using lipofectamine 2000 according to manufacturer’s instructions (Invitrogen, MA, U.S.A.). For quantification of endosomal size, EnSC cultures were fed live with anti-FLAG M1 antibody (Merck) for 20 minutes and washed before incubation with anti-mouse 488 for 20 minutes at 37 °C. Cells were washed and following ligand stimulation at 37 °C, were imaged live in phenol-free opti-MEM (ThermoFisher Scientific) using a Leica Stellaris 8 Inverted Confocal Microscope with a 63× 1.4 numerical aperture objective. Endosomal size was quantified in raw image files using ImageJ from three independent EnSC cultures. For fixed imaging, EnSCs were live fed anti-FLAG M1 antibody (Merck) for 20 minutes ± dyngo-4a pretreatment (30μM, 30 minutes), which was maintained throughout M1 and ligand treatments.

#### APPL1 co-localization

EnSCs were cultured in IBIDI 8-well chamber slides (Grafelfing, Germany). Following M1 incubation and PGE2 stimulation, the plasma membrane signal was removed by 4x PBS + 0.04M EDTA washes. Cells were immediately fixed in 4% paraformaldehyde before permeabilization (PBS Ca^2+^ + 0.2% triton x-100, 20 minutes) and blocking (PBS Ca^2+^ + 2% FCS, 30 minutes). Cells were incubated for 1 hour at RT with primary antibody for the VEE marker APPL1 (Cell Signaling Technology), before 3x PBS Ca^2+^ washes and incubated for 1 hour at RT with anti-mouse 488 ± anti-rabbit 555 secondary antibodies (ThermoFisher Scientific). Cells were imaged directly in IBIDI chambers using an Oxford Nanoimager (Oxford Nanoimaging, Oxford, U.K) in TIRF plane with a 100x 1.45 numerical aperture objective using 488 nm and 561 nm lasers. Analysis was conducted using the ImageJ JaCoP plugin, where co-localization was quantified as the average Manders Coefficient across 3 regions of 10 μM^2^ per cell using 10 cells per condition.

### RNA-sequencing

#### Cell culture

3 independent EnSC cultures were seeded into 6-well plates and transfected with GIPC, APPL1 or NT siRNA, before decidualization with PGE2/MPA for 4 days.

#### RNA isolation

RNA was extracted and purified through columns using Qiagen’s RNeasy Plus Universal Mini kit exactly as per manufacturer’s instructions.

#### RNA-Sequencing

RNA concentration and purity was assessed using the Qubit RNA BR assay, and RNA Quality was analyzed on an Agilent 2100 Bioanalyzer (Agilent Technologies, Santa Clara, CA, U.S.A>). All samples achieved a RNA Integrity Number (RIN) >8.4. Libraries were prepared using the Illumina TruSeq Stranded mRNA sample prep kit according to manufacturer’s instructions and sequencing was performed on Illumina NovaSeq 6000 (Illumina, Cambridge, U.K.) with 75pb paired-end reads at the University of Warwick Genomics Facility (Warwick, U.K.).

#### Data analysis

Transcriptomic maps of paired-end reads were generated using STAR v2.5 against the hg38 reference genome. Transcript counts were assessed by HTSeq-0.6.1, and transcripts per million were calculated. ^48^ Differential gene expression between treatments was evaluated using DEseq2-1.28.1 package in R, with significance defined as an adjusted *P*-value (q-value) of <0.05 after Bonferroni false discovery rate correction.

#### KEGG analysis

The function and pathway enrichment of differentially expressed genes was performed in the Database for Annotation, Visualization and Integrated Discovery (DAVID) Bioinformatics Resource (2021), ^49,50^ using the Kyoto Encyclopedia of Genes and Genomes (KEGG) pathway database (v108.1) to explore functional interactions among identified genes. ^51^ Statistical significance was determined using the Fisher’s exact test.

### Statistics

Unless otherwise stated, data were analyzed using GraphPad Prism (v9.1.0) (GraphPad Software Inc, CA, U.S.A.). Data are presented as either raw values or as a fold-change relative to the most informative comparator. Unpaired Student’s t-test was used to compare two groups, and one-way ANOVA with Tukey’s or Dunnett’s post-hoc test for multiple comparisons for groups of 3 or more. Sequencing analysis was performed using R (v4.1.2), ^52^ where differentially expressed genes were identified following Bonferroni’s correction for multiple comparisons. In all cases, *P*<0.05 was considered significant.

## Notes

### Competing Interest Statement

The authors have declared no competing interest.

