## Supplemental Information for "Spatial targeting of the prostaglandin receptor EP2 to very early endosomes co-ordinates PGE2-mediated cAMP signaling in decidualizing human endometrium"

**Supplemental Figure S1**

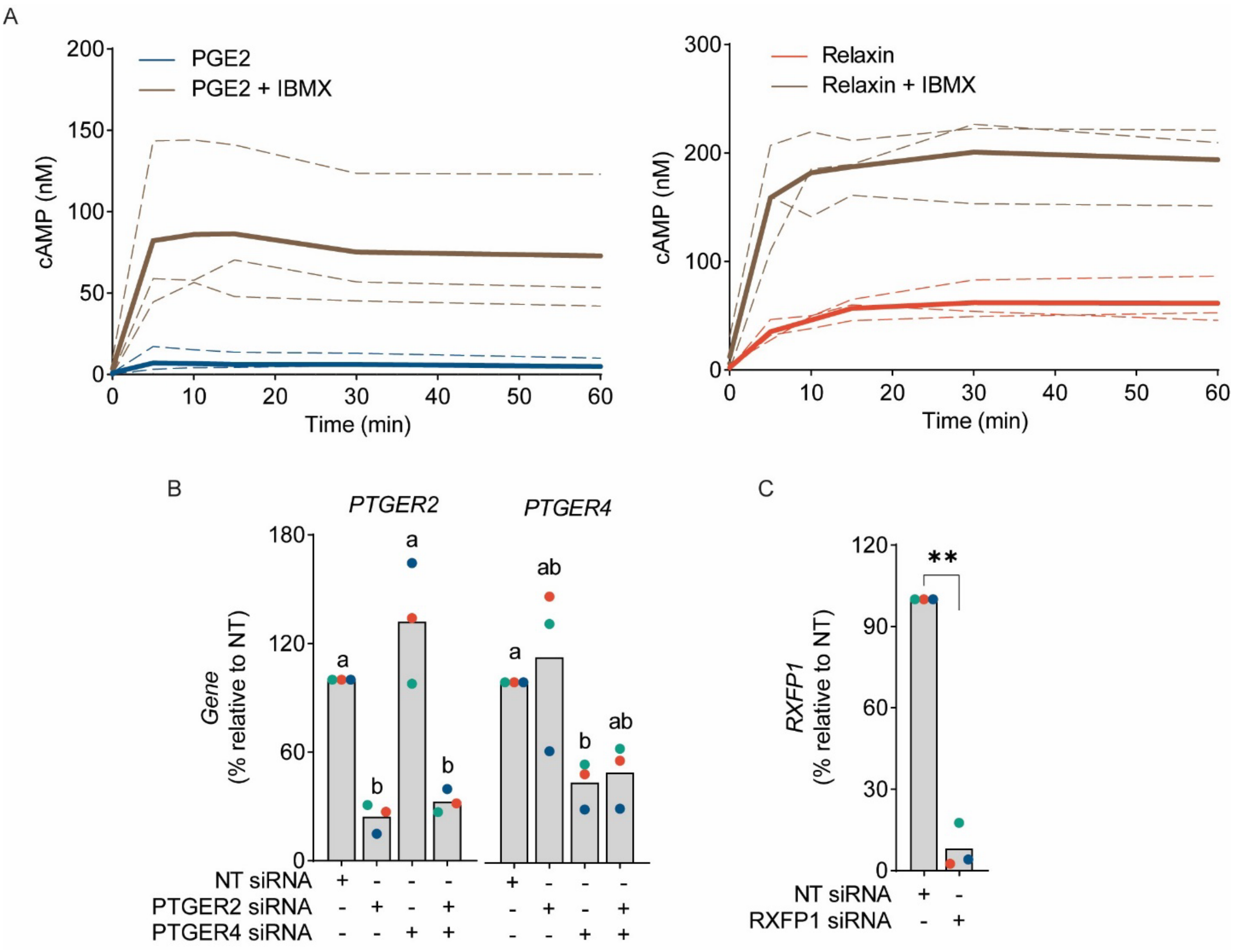


**Figure S1.** (A) Changes in cAMP induced by PGE2 or relaxin in EnSC comparing responses in the presence or absence of the phosphodiesterase inhibitor, IBMX. Plots from individual patients are represented by dashed lines with bold lines indicating mean values, n=3. (B-C) Depletion of *PTEGR2* and/or *PTGER4* (B) and *RXFP1* (C) in EnSC as determined by RTqPCR. Data points from individual patients are color-matched and shown together with bar graphs denoting mean. Different letters indicate statistical difference ($P$<0.05) from NT siRNA, stimulated cells (ANOVA and Dunnett's multiple comparison test) with ** indicating $P$<0.001 from Student's t-test, n=3.

**Supplemental Figure S2**

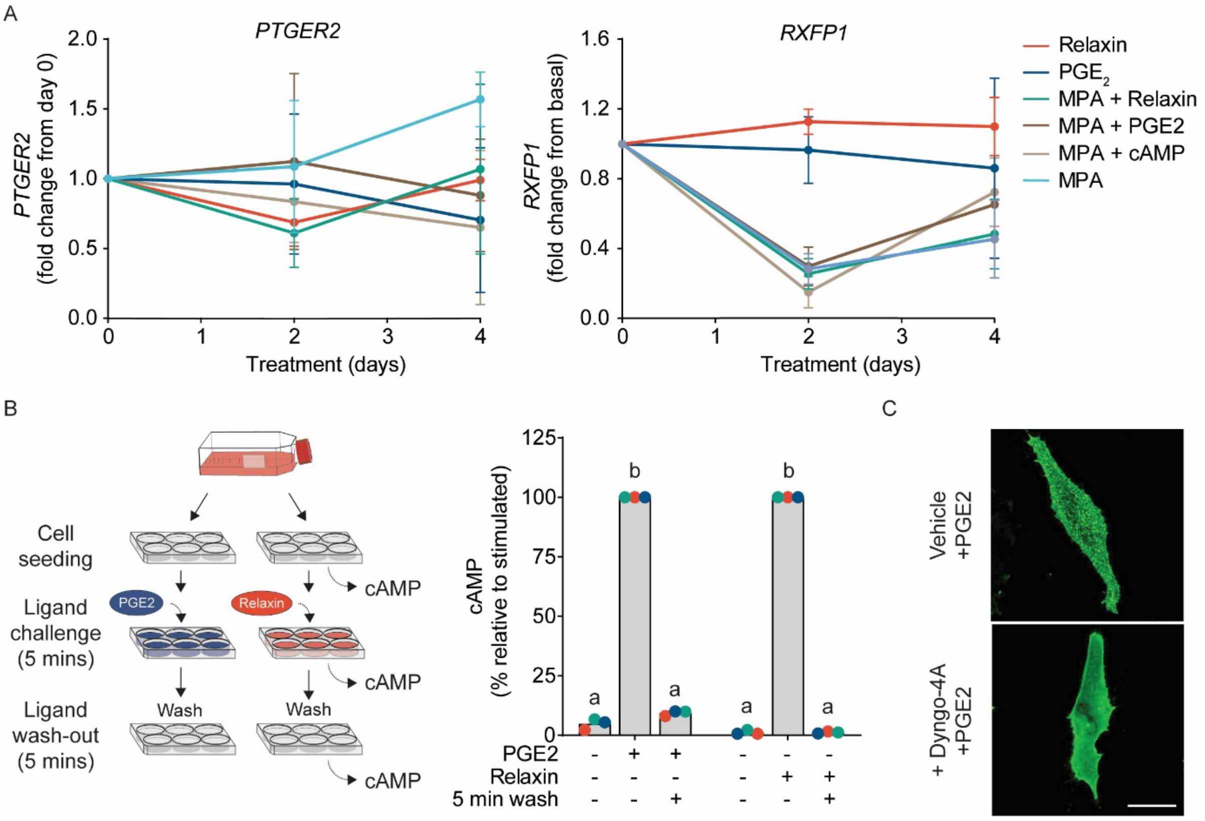


**Figure S2:** (A) RTqPCR analysis of transcripts coding EP2 (*PTGER2*) and RXFP1 (*RXFP1*) receptors following various treatments for 2 or 4 days. Data are mean ± SD, n=3. (B) Schematic representation of the experimental procedures used to assess ligand wash out (left panel). Induction of cAMP in endometrial stromal cells following 5-minute ligand wash-out (right panel). Data are shown as individual biological replicates from independent primary cultures with bars denoting mean values. Different letters indicate statistical difference ($P<0.05$) from treated cells (ANOVA and Šidák's multiple comparison test), n=3. (C) Confocal images of FLAG-Tagged EP2 receptors in a cultured EnSC pre-treated with or without the dynamin inhibitor Dyngo-4A to prevent receptor internalization. Images are representative of 3 further cultures. Scale bar = 20µm.

### Supplemental Figure S3

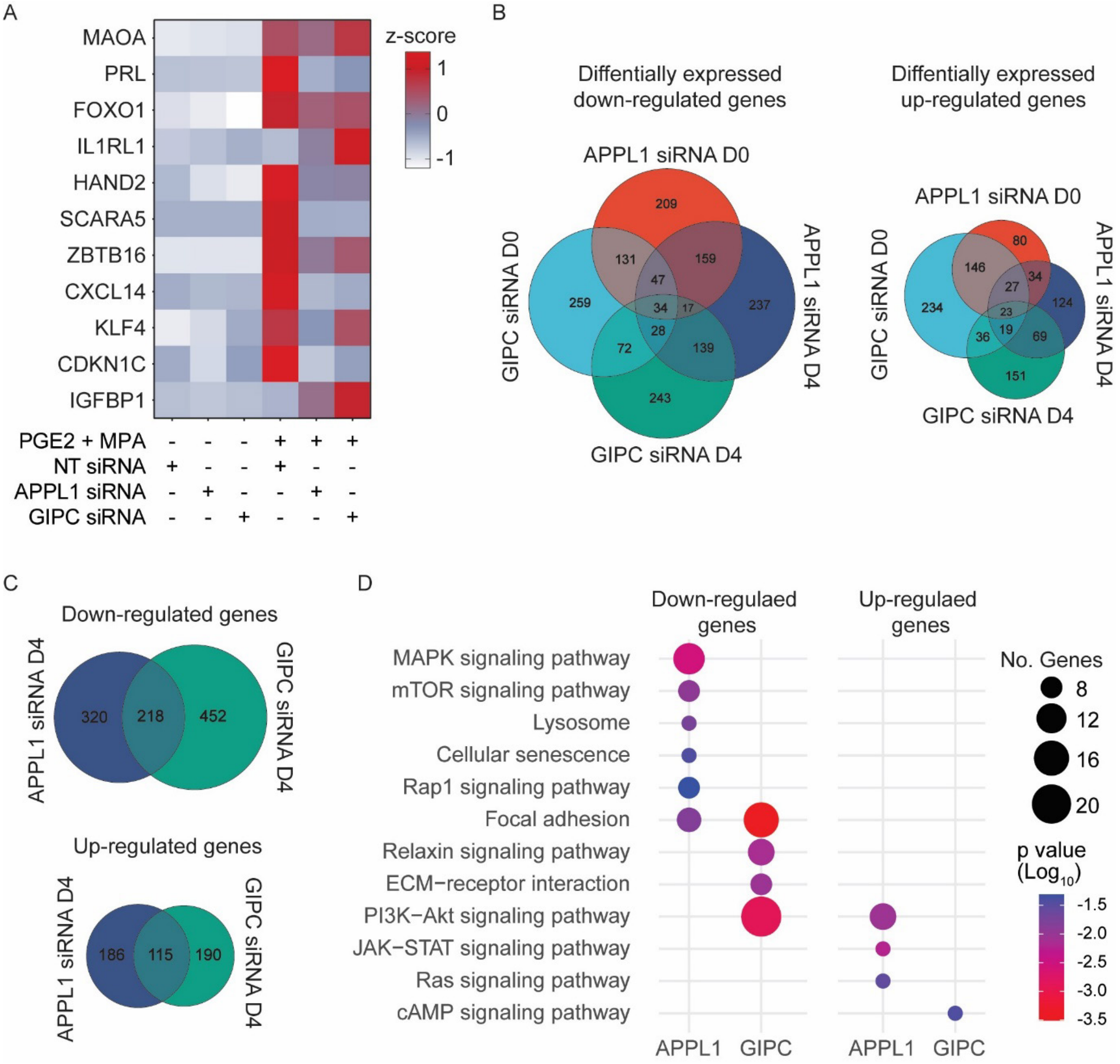


**Figure S3:** (A) Heatmap showing relative expression (z-score) of key decidual genes following depletion of APPL1 or GIPC. Red, blue and white represent high, medium and low gene expression, respectively, as indicated by the color key. (B) Venn diagram depicting the number of individual and common differentially genes (Bonferroni correction, $P<0.05$) identified comparing APPL1 and GIPC depleted EnSCs to treatment-matched NT siRNA-transfected cells. Diagrams are proportional to represent down-regulated genes (left panel) and up-regulated genes (right panel). (C) Venn diagram showing the number of unique and shared differentially genes for day-4 treated cells depleted of APPL1 or GIPC. Diagrams are proportional to represent down-regulated genes (upper panel) and up-regulated genes (lower panel. (D) Selected Kyoto Encyclopedia of Genes and Genomes (KEGG) pathway enrichment of up- and downregulated differentially expressed genes that are exclusive to APPL1 or GIPC depleted EnSC. The size of circles is relative to the number of genes in each enrichment term, and the color represents *P* value calculated as a result of enrichment degree.

**Supplemental Table S1. Patient demographics:** Endometrial biopsies used for EnSC cultures.

| Figure | n | Age | BMI | LH+ |
|---|---|---|---|---|
| Total | 46 | 36.5 (34-39.75) | 24 (22-27) | 8 (7-10) |
| 1B, 1C | 3 | 37 (36-37.5) | 28 (25-28.9) | 7 (7-9) |
| 1G, 1H, S1B, S1C | 3 | 40 (37.5-40.5) | 24 (22-28) | 8 (8-9) |
| 2A | 3 | 32 (31.5-34) | 27 (26-28) | 7 (7-7) |
| 2C | 3 | 38 (36-41) | 22.4 (21.2-24.1) | 7 (7-9.5) |
| 2D, S2B | 3 | 30 (19.5-36) | 23 (21.5-27) | 9 (8.5-10) |
| 2E | 3 | 36 (34.5-38) | 23 (22-23.5) | 9 (8-9.5) |
| 3B | 4 | 40 (37-41) | 23 (22.5-24.95) | 8 (7-9) |
| 3C | 3 | 36 (33.5-38) | 22 (21.5-28.5) | 9 (7.5-10) |
| S2C | 6 | 38 (33.5-39) | 27 (26-28) | 8 (7-8.5) |
| 4A, 4B | 3 | 36 (32-37) | 25.8 (24-25.9) | 9 (7-8) |
| 4C | 3 | 43 (38.5-43) | 27 (23.5-31.1) | 10 (10-11) |
| 5B-F, S3 | 3 | 34 (32.5-36.5) | 26 (25-28) | 7 (6.5-8) |
| 2G | 3 | 39 (37-39) | 20 (19.5-21) | 8 (8-9) |
| Supple 2A | 3 | 38 (34-39) | 22 (21.5-23) | 7 (6.5-8) |

All data are median (interquartile range, Q1-Q3).
LH+: Days since the pre-ovulatory Luteinizing Hormone (LH) surge

**Supplemental Table S2.**

| NT siRNA Day 4 vs APPL1 siRNA Day 4 | | | | |
|---|---|---|---|---|
| **GO Term (KEGG Pathway)** | **No. of genes** | **%** | **Fold Enrichment** | ***P* value** |
| Focal adhesion | 31 | 3.2 | 2.8 | 5.80E-07 |
| PI3K-Akt signaling pathway | 42 | 4.4 | 2.2 | 3.90E-06 |
| Pathways in cancer | 55 | 5.7 | 1.9 | 6.20E-06 |
| Proteoglycans in cancer | 28 | 2.9 | 2.5 | 1.90E-05 |
| Protein digestion and absorption | 18 | 1.9 | 3.2 | 4.00E-05 |
| Small cell lung cancer | 16 | 1.7 | 3.2 | 1.30E-04 |
| ECM-receptor interaction | 15 | 1.6 | 3.1 | 3.20E-04 |
| Human papillomavirus infection | 33 | 3.4 | 1.8 | 1.20E-03 |
| cAMP signaling pathway | 24 | 2.5 | 1.9 | 2.90E-03 |
| Arrhythmogenic right ventricular cardiomyopathy | 12 | 1.3 | 2.8 | 3.00E-03 |
| Hypertrophic cardiomyopathy | 13 | 1.4 | 2.6 | 3.60E-03 |
| Steroid biosynthesis | 6 | 0.6 | 5.4 | 3.80E-03 |
| Dilated cardiomyopathy | 13 | 1.4 | 2.5 | 6.10E-03 |
| Toxoplasmosis | 14 | 1.5 | 2.3 | 8.20E-03 |
| AGE-RAGE signaling pathway in diabetic complications | 13 | 1.4 | 2.4 | 8.40E-03 |
| Amoebiasis | 13 | 1.4 | 2.3 | 9.80E-03 |
| Hepatocellular carcinoma | 18 | 1.9 | 1.9 | 1.10E-02 |
| EGFR tyrosine kinase inhibitor resistance | 11 | 1.1 | 2.5 | 1.10E-02 |
| Relaxin signaling pathway | 15 | 1.6 | 2.1 | 1.10E-02 |
| Rap1 signaling pathway | 21 | 2.2 | 1.8 | 1.10E-02 |
| MAPK signaling pathway | 27 | 2.8 | 1.6 | 1.50E-02 |
| Platinum drug resistance | 10 | 1 | 2.5 | 1.80E-02 |
| Complement and coagulation cascades | 11 | 1.1 | 2.3 | 1.90E-02 |
| JAK-STAT signaling pathway | 17 | 1.8 | 1.9 | 2.00E-02 |
| Fluid shear stress and atherosclerosis | 15 | 1.6 | 2 | 2.00E-02 |
| Calcium signaling pathway | 23 | 2.4 | 1.7 | 2.20E-02 |
| Axon guidance | 18 | 1.9 | 1.8 | 2.20E-02 |
| Pancreatic cancer | 10 | 1 | 2.4 | 2.30E-02 |
| Chagas disease | 12 | 1.3 | 2.1 | 2.40E-02 |
| Glycosaminoglycan | 5 | 0.5 | 4.3 | 2.60E-02 |
| Growth hormone synthesis, secretion and action | 13 | 1.4 | 2 | 3.20E-02 |
| Ras signaling pathway | 21 | 2.2 | 1.6 | 3.50E-02 |
| Platelet activation | 13 | 1.4 | 1.9 | 3.90E-02 |
| Circadian entrainment | 11 | 1.1 | 2.1 | 4.00E-02 |
| Other types of O-glycan biosynthesis | 7 | 0.7 | 2.7 | 4.20E-02 |
| ErbB signaling pathway | 10 | 1 | 2.1 | 4.30E-02 |
| Melanoma | 9 | 0.9 | 2.3 | 4.30E-02 |
| Oxytocin signaling pathway | 15 | 1.6 | 1.8 | 4.30E-02 |
| Cushing syndrome | 15 | 1.6 | 1.8 | 4.50E-02 |

**Table S2**. Full list of KEGG terms from differentially expressed genes comparing APPL1 and NT siRNA-transfected EnSCs stimulated with PGE2 and MPA for 4 days.

### Supplemental Table S3.

| NT siRNA Day 4 vs GIPC siRNA Day 4 | | | | |
|---|---|---|---|---|
| **GO Term (KEGG Pathway)** | **No. of genes** | **%** | **Fold Enrichment** | ***P* value** |
| cAMP signaling pathway | 26 | 3.1 | 2.6 | 2.40E-05 |
| Arrhythmogenic right ventricular cardiomyopathy | 14 | 1.7 | 4 | 3.30E-05 |
| Parathyroid hormone synthesis, secretion and action | 16 | 1.9 | 3.4 | 7.10E-05 |
| Focal adhesion | 23 | 2.8 | 2.5 | 1.10E-04 |
| Cushing syndrome | 19 | 2.3 | 2.7 | 1.90E-04 |
| Vascular smooth muscle contraction | 17 | 2.1 | 2.8 | 3.10E-04 |
| Dilated cardiomyopathy | 14 | 1.7 | 3.2 | 3.30E-04 |
| MAPK signaling pathway | 28 | 3.4 | 2.1 | 4.70E-04 |
| cGMP-PKG signaling pathway | 19 | 2.3 | 2.5 | 4.80E-04 |
| Adrenergic signaling in cardiomyocytes | 18 | 2.2 | 2.6 | 5.20E-04 |
| Oxytocin signaling pathway | 18 | 2.2 | 2.6 | 5.20E-04 |
| Cortisol synthesis and secretion | 11 | 1.3 | 3.8 | 5.80E-04 |
| Calcium signaling pathway | 24 | 2.9 | 2.1 | 9.80E-04 |
| Axon guidance | 19 | 2.3 | 2.3 | 1.30E-03 |
| Regulation of actin cytoskeleton | 22 | 2.7 | 2.1 | 1.40E-03 |
| Hypertrophic cardiomyopathy | 12 | 1.5 | 3 | 2.20E-03 |
| Long-term potentiation | 10 | 1.2 | 3.3 | 2.90E-03 |
| mTOR signaling pathway | 16 | 1.9 | 2.3 | 4.30E-03 |
| Toxoplasmosis | 13 | 1.6 | 2.6 | 4.30E-03 |
| Proteoglycans in cancer | 19 | 2.3 | 2.1 | 4.90E-03 |
| TNF signaling pathway | 13 | 1.6 | 2.5 | 5.00E-03 |
| Rap1 signaling pathway | 19 | 2.3 | 2 | 6.30E-03 |
| Cholesterol metabolism | 8 | 1 | 3.5 | 7.30E-03 |
| Pathways in cancer | 37 | 4.5 | 1.5 | 8.20E-03 |
| Wnt signaling pathway | 16 | 1.9 | 2.1 | 9.80E-03 |
| Aldosterone synthesis and secretion | 11 | 1.3 | 2.5 | 1.20E-02 |
| Human papillomavirus infection | 25 | 3 | 1.7 | 1.40E-02 |
| Shigellosis | 20 | 2.4 | 1.8 | 1.50E-02 |
| Long-term depression | 8 | 1 | 3 | 1.70E-02 |
| Growth hormone synthesis, secretion and action | 12 | 1.5 | 2.2 | 1.90E-02 |
| Gastric acid secretion | 9 | 1.1 | 2.6 | 2.00E-02 |
| Adherens junction | 10 | 1.2 | 2.4 | 2.30E-02 |
| Pathways of neurodegeneration - multiple diseases | 32 | 3.9 | 1.5 | 2.30E-02 |
| Human cytomegalovirus infection | 18 | 2.2 | 1.8 | 2.50E-02 |
| PI3K-Akt signaling pathway | 25 | 3 | 1.6 | 2.80E-02 |
| Human immunodeficiency virus 1 infection | 17 | 2.1 | 1.8 | 2.90E-02 |
| Circadian entrainment | 10 | 1.2 | 2.3 | 3.00E-02 |
| Glutamatergic synapse | 11 | 1.3 | 2.1 | 3.40E-02 |
| Serotonergic synapse | 11 | 1.3 | 2.1 | 3.40E-02 |
| Melanogenesis | 10 | 1.2 | 2.2 | 3.70E-02 |
| Hepatocellular carcinoma | 14 | 1.7 | 1.9 | 3.90E-02 |
| Progesterone-mediated oocyte maturation | 10 | 1.2 | 2.2 | 3.90E-02 |
| Cardiac muscle contraction | 9 | 1.1 | 2.3 | 4.10E-02 |
| Human T-cell leukemia virus 1 infection | 17 | 2.1 | 1.7 | 4.20E-02 |
| ECM-receptor interaction | 9 | 1.1 | 2.2 | 4.60E-02 |
| Estrogen signaling pathway | 12 | 1.5 | 1.9 | 4.60E-02 |
| Apelin signaling pathway | 12 | 1.5 | 1.9 | 4.80E-02 |

**Table S3:** Full list of KEGG terms from differentially expressed genes comparing GIPC and NT siRNA-transfected EnSCs stimulated with PGE2 and MPA for 4 days.

### Supplemental Table S4.

| NT siRNA Day 0 vs APPL1 siRNA Day 0 | | | | |
|---|---|---|---|---|
| **GO Term (KEGG Pathway)** | **No. of genes** | **%** | **Fold Enrichment** | ***P* value** |
| Focal Adhesion | 21 | 2.3 | 2.1 | 2.40E-03 |
| Complement and coagulation cascades | 12 | 1.3 | 2.8 | 3.20E-03 |
| Protein digestion and absorption | 13 | 1.4 | 2.6 | 4.60E-03 |
| Insulin resistance | 13 | 1.4 | 2.4 | 6.70E-03 |
| Human immunodeficiency virus 1 infection | 20 | 2.2 | 1.9 | 8.30E-03 |
| Regulation of actin cytoskeleton | 21 | 2.3 | 1.9 | 9.10E-03 |
| TNF signaling pathway | 13 | 1.4 | 2.3 | 1.00E-02 |
| Relaxin signaling pathway | 14 | 1.5 | 2.2 | 1.10E-02 |
| Proteoglycans in cancer | 19 | 2.1 | 1.9 | 1.20E-02 |
| Arrhythmogenic right ventricular cardiomyopathy | 10 | 1.1 | 2.6 | 1.30E-02 |
| Hypertrophic cardiomyopathy | 11 | 1.2 | 2.5 | 1.30E-02 |
| Growth hormone synthesis, secretion and action | 13 | 1.4 | 2.2 | 1.50E-02 |
| Human cytomegalovirus infection | 20 | 2.2 | 1.8 | 1.50E-02 |
| Yersinia infection | 14 | 1.5 | 2.1 | 1.70E-02 |
| Fluid shear stress and atherosclerosis | 14 | 1.5 | 2 | 1.90E-02 |
| Circadian entrainment | 11 | 1.2 | 2.3 | 2.10E-02 |
| Epithelial cell signaling in Helicobacter pylori infection | 9 | 1 | 2.6 | 2.10E-02 |
| Human papillomavirus infection | 26 | 2.9 | 1.6 | 2.20E-02 |
| Hippo signaling pathway | 15 | 1.7 | 1.9 | 2.30E-02 |
| Type II diabetes mellitus | 7 | 0.8 | 3.1 | 2.40E-02 |
| AGE-RAGE signaling pathway in diabetic complications | 11 | 1.2 | 2.2 | 2.50E-02 |
| MAPK signaling pathway | 24 | 2.6 | 1.6 | 2.50E-02 |
| PI3K-Akt signaling pathway | 27 | 3 | 1.5 | 2.70E-02 |
| Chagas disease | 11 | 1.2 | 2.2 | 2.80E-02 |
| Hepatitis B | 15 | 1.7 | 1.9 | 2.90E-02 |
| Valine, leucine and isoleucine degradation | 7 | 0.8 | 3 | 2.90E-02 |
| Rap1 signaling pathway | 18 | 2 | 1.7 | 3.00E-02 |
| Longevity regulating pathway | 10 | 1.1 | 2.3 | 3.10E-02 |
| Chronic myeloid leukemia | 9 | 1 | 2.4 | 3.30E-02 |
| Axon guidance | 16 | 1.8 | 1.8 | 3.50E-02 |
| cGMP-PKG signaling pathway | 15 | 1.7 | 1.8 | 3.60E-02 |
| Insulin signaling pathway | 13 | 1.4 | 1.9 | 3.70E-02 |
| Phagosome | 14 | 1.5 | 1.9 | 3.70E-02 |
| Salmonella infection | 20 | 2.2 | 1.6 | 3.80E-02 |
| Oxytocin signaling pathway | 14 | 1.5 | 1.8 | 4.00E-02 |
| TGF-beta signaling pathway | 11 | 1.2 | 2.1 | 4.00E-02 |
| Endocytosis | 20 | 2.2 | 1.6 | 4.10E-02 |
| Pathways of neurodegeneration - multiple diseases | 33 | 3.6 | 1.4 | 4.40E-02 |
| Autophagy - animal | 13 | 1.4 | 1.9 | 4.50E-02 |
| N-Glycan biosynthesis | 7 | 0.8 | 2.7 | 4.50E-02 |
| Dilated cardiomyopathy | 10 | 1.1 | 2.1 | 4.60E-02 |
| Human T-cell leukemia virus 1 infection | 18 | 2 | 1.6 | 4.70E-02 |
| Protein processing in endoplasmic reticulum | 15 | 1.7 | 1.7 | 4.80E-02 |
| Toxoplasmosis | 11 | 1.2 | 2 | 4.90E-02 |

**Table S4:** Full list of KEGG terms from differentially expressed genes comparing unstimulated (Day 0) APPL1 and NT siRNA-transfected EnSCs.

### Supplemental Table S5.

| NT siRNA Day 0 vs GIPC siRNA Day 0 | | | | |
|---|---|---|---|---|
| **GO Term (KEGG Pathway)** | **No. of genes** | **%** | **Fold Enrichment** | ***P* value** |
| Proteoglycans in cancer | 30 | 2.9 | 2.5 | 1.00E-05 |
| Vascular smooth muscle contraction | 21 | 2 | 2.6 | 1.10E-04 |
| Wnt signaling pathway | 23 | 2.2 | 2.3 | 4.70E-04 |
| TGF-beta signaling pathway | 17 | 1.6 | 2.6 | 5.80E-04 |
| Pathways in cancer | 51 | 4.9 | 1.6 | 6.70E-04 |
| Arrhythmogenic right ventricular cardiomyopathy | 13 | 1.2 | 2.8 | 1.80E-03 |
| Hippo signaling pathway | 20 | 1.9 | 2.1 | 2.30E-03 |
| IL-17 signaling pathway | 14 | 1.3 | 2.5 | 3.50E-03 |
| Spinocerebellar ataxia | 18 | 1.7 | 2.1 | 4.60E-03 |
| Breast cancer | 18 | 1.7 | 2.1 | 6.10E-03 |
| Hypertrophic cardiomyopathy | 13 | 1.2 | 2.4 | 6.60E-03 |
| MAPK signaling pathway | 30 | 2.9 | 1.7 | 6.80E-03 |
| Renin secretion | 11 | 1 | 2.7 | 7.10E-03 |
| NF-kappa B signaling pathway | 14 | 1.3 | 2.3 | 8.30E-03 |
| Focal adhesion | 22 | 2.1 | 1.8 | 8.90E-03 |
| Regulation of actin cytoskeleton | 24 | 2.3 | 1.8 | 9.00E-03 |
| Shigellosis | 25 | 2.4 | 1.7 | 1.10E-02 |
| NOD-like receptor signaling pathway | 20 | 1.9 | 1.8 | 1.40E-02 |
| Calcium signaling pathway | 25 | 2.4 | 1.7 | 1.50E-02 |
| Gap junction | 12 | 1.1 | 2.3 | 1.50E-02 |
| Yersinia infection | 16 | 1.5 | 2 | 1.50E-02 |
| Estrogen signaling pathway | 16 | 1.5 | 2 | 1.60E-02 |
| TNF signaling pathway | 14 | 1.3 | 2.1 | 1.70E-02 |
| Human papillomavirus infection | 30 | 2.9 | 1.5 | 2.20E-02 |
| Pathogenic Escherichia coli infection | 20 | 1.9 | 1.7 | 2.40E-02 |
| Dilated cardiomyopathy | 12 | 1.1 | 2.1 | 2.60E-02 |
| Sphingolipid signaling pathway | 14 | 1.3 | 1.9 | 2.70E-02 |
| Pertussis | 10 | 1 | 2.2 | 3.50E-02 |
| Pathways of neurodegeneration - multiple diseases | 39 | 3.7 | 1.4 | 3.60E-02 |
| Sphingolipid metabolism | 8 | 0.8 | 2.5 | 3.60E-02 |
| JAK-STAT signaling pathway | 17 | 1.6 | 1.7 | 3.70E-02 |
| Leishmaniasis | 10 | 1 | 2.2 | 3.80E-02 |
| cGMP-PKG signaling pathway | 17 | 1.6 | 1.7 | 3.90E-02 |
| Salmonella infection | 23 | 2.2 | 1.6 | 3.90E-02 |
| Axon guidance | 18 | 1.7 | 1.7 | 4.20E-02 |
| Rap1 signaling pathway | 20 | 1.9 | 1.6 | 4.30E-02 |
| Tight junction | 17 | 1.6 | 1.7 | 4.30E-02 |
| Cytokine-cytokine receptor interaction | 26 | 2.5 | 1.5 | 4.80E-02 |

**Table S5:** Full list of KEGG terms from differentially expressed genes comparing unstimulated (Day 0) GIPC and NT siRNA-transfected EnSCs.
